# m6ASeqTools: An R toolkit for post-processing m6A sites detected by m6Anet

**DOI:** 10.1101/2025.08.15.670588

**Authors:** Hanna Lee, Denalda Gashi, Syeda Maheen Batool, Ana K. Escobedo, Bob S. Carter, Leonora Balaj

## Abstract

m6ASeqTools is an R package designed to streamline the post-processing and interpretation of site-level m6A predictions from m6Anet. It provides descriptive summaries of m6A distribution across transcripts, genes, biotypes, and transcript regions, and enables condition comparisons through a calculated weighted modification ratio. By integrating differential gene expression data, the package links methylation changes with expression differences, providing biotype-specific and region-specific insights into how m6A localization patterns relate to transcriptional regulation.

**Availability and implementation:** m6ASeqTools is freely available at https://github.com/hannalee809/m6ASeqTools

**Supplementary information:** Supplementary data are available at Bioinformatics online.

## Introduction

N6-methyladenosine (m6A) is an abundant RNA modification that regulates mRNA stability, splicing, and translation (Lang et al. 2025; Tang, Gupta, and Guo 2022; Zhou et al. 2022). Aberrant m6A regulation has been associated with various diseases, including cancer and neurological disorders (Pinello, Sun, and Wong 2018). Given the central role in m6A machinery in disease pathogenesis, comprehensive transcriptome-wide profiling of m6A modifications is critical for uncovering new biological mechanisms and potential therapeutic targets.

Recent advancements in direct RNA sequencing technologies, particularly those from Oxford Nanopore Technologies, now enable the precise, single-nucleotide resolution mapping of m6A sites. This capability has further fueled the emergence of sophisticated prediction models, such as m6Anet, a deep learning-based tool that provides high-resolution, transcriptome-wide identification of m6A modifications (Hendra et al. 2022). The output from m6Anet offers detailed information on identified m6A sites, including transcript coordinates, kmer modified, modification probability and modification ratio.

While m6Anet provides valuable site-level information, the need for robust downstream analysis tools to interpret and contextualize these predictions remains critical. To address this gap, we present m6ASeqTools, an R package specifically designed for the post-processing of m6Anet data. m6ASeqTools enables researchers to derive important descriptive statistics of m6A modifications and integrates reference genome annotations to contextualize these modifications at the transcript and gene levels. Furthermore, recognizing the importance of comparative studies, m6ASeqTools introduces a transcript-level methylation metric, facilitating meaningful comparisons of m6A levels across experimental conditions. To this end, the package integrates gene expression analysis and provides summaries of transcript biotypes and region-specific modification patterns, offering deeper insights into the functional relevance of m6A.

### Features and Implementation

m6ASeqTools utilizes the *data.site_proba.csv* file generated by m6Anet, which provides site-level information on m6A modifications. This includes transcript position, transcript biotype, modified k-mer sequence, the predicted probability of modification, and the estimated modification ratio for each site. To enable transcript-level summarization, users must also provide a corresponding GTF annotation file (e.g. form GENCODE or Ensembl) to extract transcript structure information and calculate region-specific and total transcript lengths. Additionally, m6ASeqTools supports integration with gene expression data for joint analyses of RNA methylation and expression. Users can provide differential expression results, such as the output from DESeq2 (Love, Huber, and Anders 2014), containing log2 fold change and adjusted p-values for the comparison of interest.

m6ASeqTools provides two main analytical modules: (1) descriptive statistics of m6A methylation patterns, and (2) comparative analysis across experimental groups. Starting from m6Anet’s data.site_proba.csv output, the package computes and visualizes key metrics such as the distribution of modified sites, transcripts, and genes, the frequency of transcript biotypes, k-mer enrichment, and the length distribution of modified transcripts. These statistics are organized into an HTML summary report, with all tables and plots rendered using ggplot2.

Chromosome-level distributions of modified genes are generated using org.Hs.eg.db (Carlson 2017) as the reference genome annotation. To compute the relative positions of m6A sites along the transcript regions (5’UTR, CDS, 3’UTR), this package utilizes the user-supplied GTF annotation file and visualizes the spatial distribution using kernel density plots.

For group-level comparisons, m6ASeqTools identifies common and unique m6A modified sites, transcripts, and genes across any number of user-defined sample groups. To quantify transcript-level methylation, a weighted modification ratio is calculated for each transcript as:

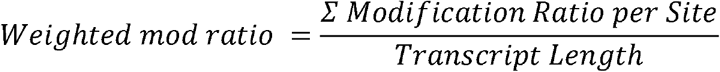

Additionally, the package offers integration with differential gene expression (DGE) analysis. When DESeq2 results are provided, users can assess the relationship between differential transcript-level methylation and gene expression changes, including visualization of log2 fold changes from both analyses on a single plot, enabling exploration of functional consequences of m6A modifications. To further interpret these relationships, m6ASeqTools classifies transcripts into six functional categories based on combined methylation and expression profiles: transcripts that are hypermethylated and have an upregulated gene expression in a group, hypermethylated with downregulated gene expression, and hypermethylated with no significant change in gene expression. For each category, the package generates summaries of transcript biotype distributions and m6A site localizations across transcript regions, providing a structured overview of the biological context associated with differential methylation. The package includes two automated functions designed to streamline statistical analysis. m6ASeqTools_part1() generates descriptive statistics from the m6Anet site probability output. m6ASeqTools_part2() uses the results from the first function to perform comparative analysis between two groups, integrating transcript methylation and gene expression to generate further insights into the functional consequences of m6A modifications, providing quantitative measures to evaluate differences in methylation patterns, transcript biotypes, and regional distributions between conditions.

### Application

The analytical workflow implemented here was adapted from the approaches described in Batool et al. (2024) and Batool et al. (2024) and integrated into m6ASeqTools. The package was applied to the glioma cell line dataset as part of our cell lines project, which is currently under review. We utilized m6ASeqTools to analyze long-read RNA sequencing data derived from the human glioma cell line Gli36 (Jones et al., 2019). In this application, we focused on the knockdown of IGF2BP2, an m6A reader implicated in cancer progression through the regulation of m6A-modified miRNAs, lncRNAs, and other transcripts (Wang, Chen, and Qiang, 2021). Our goal was to investigate transcriptome-wide changes in m6A methylation and gene expression resulting from IGF2BP2 knockdown compared to an untreated Naive condition. For each condition, cell RNA was sequenced with three technical replicates per sample. Basecalling was performed using Dorado from Oxford Nanopore Technologies, followed by alignment to the human transcriptome using minimap2 (Li, 2018). Resultant BAM files, along with associated FAST5, FASTQ, and reference transcriptome files, were processed using m6Anet. Replicates were pooled using m6Anet’s native pooling functionality, producing four output files *data.site_proba.csv* for subsequent analysis.

Application of m6ASeqTools reveals several notable observations (**Fig. 1**). The Naive condition exhibited the highest number of high-confidence m6A-modified sites (probability of modification > 0.9), transcripts, and genes compared to IGF2BP2 KD (**Supplementary Fig. 1a**). Although the majority of modified transcripts had a single modified site, the Naive condition had up to 13 modified sites per transcript and IGF2BP2 KD had up to 7 (**Fig. 1c**). Modified transcripts per gene were similar across both groups (Naive: up to 17, IGF2BP2 KD: up to 19) (**Supplementary Fig. 1b**).

**Figure 1.**
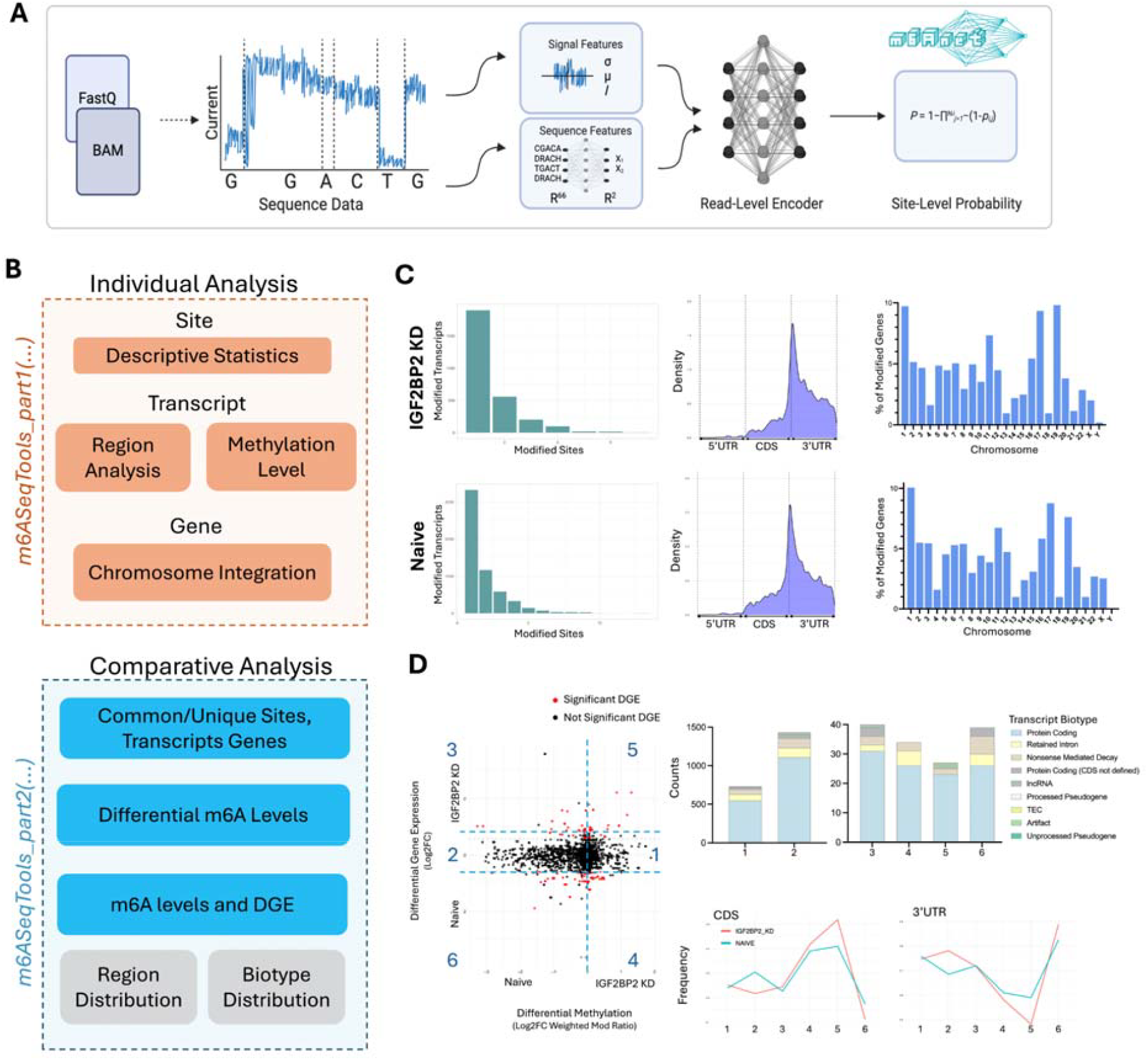
Overview of the m6ASeqTools pipeline. **(a)** Signal and sequence features from m6Anet are used to predict site-level m6A probabilities, generating a data.site_proba.csv file that serves as input for m6ASeqTools. **(b)** The pipeline includes two modules: Part 1 (orange) produces descriptive statistics of m6A methylation patterns for individual samples, while Part 2 (blue) performs comparative analyses between groups. **(c)** Example part 1 outputs for IGF2BP2 knockdown and Naïve conditions include distributions of single- and multi-site methylated transcripts, m6A site localization along transcript regions, chromosomal distributions of modified genes, **(d)** Example part 2 outputs comparing IGF2BP2 KD vs Naïve include scatter plots integrating differential methylation with gene expression. Numbered clusters from the scatter plot are further analyzed for transcript biotype and region composition.

Motif analysis demonstrated consistent enrichment of canonical DRACH motifs across conditions, with GGACT, GAACA, and GAACT identified as the most frequently modified sequences (**Supplementary Fig. 1c)**. The majority of modified transcripts were protein-coding (Naive: 73.7%; IGF2BP2 KD: 75.5%), followed by nonsense-mediated decay (Naive: 11.0%; IGF2BP2 KD: 9.5%) and retained intron transcripts (Naive: 10.1%; IGF2BP2 KD: 9.5%). Compared to the Naive condition, IGF2BP2 KD showed a slightly higher proportion of modified protein-coding transcripts, whereas the Naive condition had a greater proportion of non-coding transcripts (**Supplementary Fig. 1c**). Distribution of the m6A sites along the transcripts highlighted an enrichment within the 3’UTR with a peak at the start of the 3’UTR (**Fig. 1c**). Notably, the Naive condition exhibited a slighter higher density of m6A modifications within the CDS compared to experimental conditions, whereas IGF2BP2 KD demonstrated a shift towards increased 3’UTR methylation. Analysis of transcript lengths indicated that the majority of modified transcripts across all groups fell within the 5001–10,000 nucleotide range (**Supplementary Fig. 1c**). Similar distribution of modified transcript lengths were observed, with 84% of all modified transcripts in the range of 1001-5000bp in both conditions. Chromosome-level distribution of modified genes was remarkably similar across conditions with chromosomes 1, 17 and 19 most frequently modified **(Fig. 1c**). However, IGF2BP2 had higher modification frequency in chromosome 1 (9.7%) and 19 (9.8%), followed by chromosome 1 (9.3%). Naive had higher modification frequency in chromosome 1 (10.1%), followed by 17 (8.8%) and 19 (7.6%; **Fig. 1c**).

Part two of m6ASeqTools revealed distinct relationships between transcript methylation and gene expression. The package identified 3,327 sites, 2,325 transcripts, and 907 genes that were commonly modified in both conditions (**Supplementary Fig. 1d**). Across these, the Naive condition had nearly twice as many hypermethylated transcripts (n=1534) as IGF2BP2 KD (n=791; **Fig. 1d**). Of the 1,534 Naive hypermethylated transcripts, only 10 genes were significantly upregulated relative to the knockdown condition, whereas 11 genes from the 791 IGF2BP2 KD hypermethylated transcripts were significantly upregulated compared to Naive.

Transcript biotype composition also differed between groups. In the IGF2BP2 KD condition, hypermethylated transcripts with upregulated expression primarily consisted of protein coding (85.1%) with a smaller fraction of nonsense mediated decay transcripts (7.4%; **Fig. 1d**). In contrast, Naive hypermethylated and upregulated genes displayed a more diverse profile: protein coding (66.7%), retained Intron (10.3%), nonsense-mediated decay (15.4%), and protein coding with undefined CDS (7.7%; **Fig. 1d**). Among the hypermethylated transcripts with no significant changes in gene expression, Naive had a higher frequency of hypermethylated protein coding transcripts (77.3% vs 74.8%), while IGF2BP2 KD contained more retained intron transcripts (11.1% vs 8.5%, **Fig. 1d**).

Regional localization patterns provided additional insight. Hypermethylated transcripts with upregulated expression in IGF2BP2 KD were modified preferentially within the CDS region, whereas Naive exhibited enrichment almost exclusively in the 3′UTR (**Fig. 1d)**. For hypermethylated transcripts without significant expression changes, both conditions displayed a similar distribution of modifications between the CDS and 3′UTR.

Overall, these results provide preliminary evidence that IGF2BP2 knockdown reshapes the m6A landscape. We observed differences in the number of modified sites, transcripts, and genes, as well as transcript biotype composition and the regional localization of m6A marks. Such patterns, particularly the shift from CDS to 3’UTR enrichment and changes in the prevalence of protein coding versus non-coding transcripts, highlight the nuanced regulatory roles of IGF2BP2 in shaping the epitranscriptomic landscape.

## Conclusion

Recent advances in m6A prediction tools, such as m6Anet, has significantly improved the robustness and accuracy of site-level m6A modification detection. Building upon these developments, m6ASeqTools complements these predictions by providing streamlined post-processing and biological interpretation, connecting site-level modification data to transcript- and gene-level insights. By summarizing distributions, regional mapping, and integrating gene expression analysis, the package facilitates a deeper understanding of the functional roles of m6A. Future developments will focus on expanding m6ASeqTools to include additional downstream analysis modules and to support outputs from other m6A detection platforms.

## Data availability Statement

The data underlying this article will be shared on reasonable request to the corresponding author.

