## supplemental figure for "m6ASeqTools: An R toolkit for post-processing m6A sites detected by m6Anet"

**Supplementary Figure 1**

**A**

| Condition | Total Sites | Total Transcripts | Total Genes | >.90 Sites | >.90 Transcripts | >.90 Genes |
|---|---|---|---|---|---|---|
| IGF2BP2 KD | 257808 | 17627 | 4976 | 4228 | 2796 | 1067 |
| Naive | 526317 | 24438 | 6773 | 10605 | 5710 | 2047 |

**B**

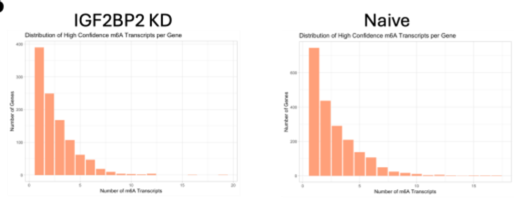


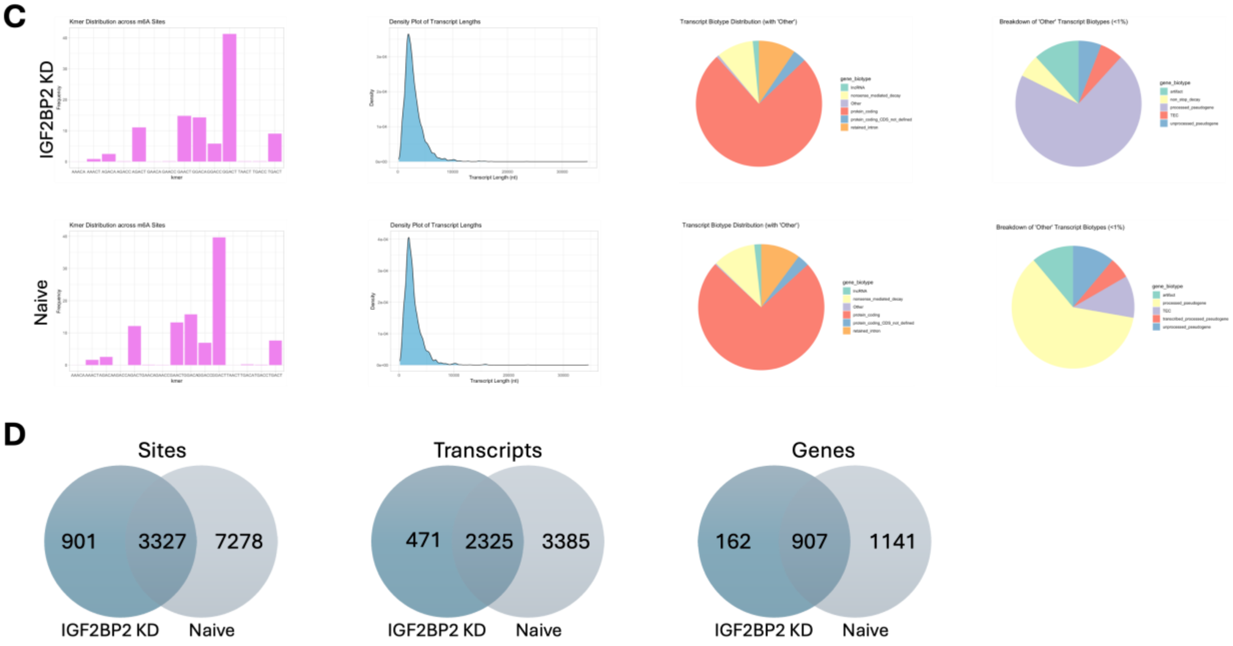


**Supplementary Fig. 1. m6ASeqTools Output for IGF2BP2 KD and Naïve conditions. (a)** Table of total and high-confidence (probability_modified > .9) m6A sites, transcripts and genes. (**b**) Distribution of the number of genes with associated counts of modified transcripts per gene. **(c)** Descriptive statistics of m6A modifications for IGF2BP2 (top) and Naïve (bottom), including kmer distribution, modified transcript length and biotype distribution. Pie chart on the left shows the frequency of modified transcript biotypes with "Other" category, which is then broken down in pie chart on the right. **(d)** Venn Diagrams showing the distribution of common and unique sites (left), transcripts (middle) and genes (right) between the two conditions.
